# Size-dependent harvest mortality indirectly affects boldness, feeding rate, and behaviour-linked gene expression in a decade-long selection experiment on guppies

**DOI:** 10.1101/2023.11.28.569017

**Authors:** Beatriz Diaz Pauli, Valentina Tronci, Ana S. Gomes, Mikko Heino

## Abstract

Fisheries-induced mortality is size-selective, commonly targeting large individuals, which leads to evolution towards smaller size and early maturation. However, little is known on whether behaviour is affected. Here we aimed at testing whether size-dependent harvest indirectly affects behavioural traits that might have ecological consequences. Specifically, we assessed feeding rate – which affects prey abundance –, boldness – which determines a fish vulnerability to predators –, and sociability – which determines how a fish interacts with conspecifics and ultimately affects foraging and predation avoidance. In addition, we tested whether the differences in behaviour were associated to differences in selected key genes expression, to understand its molecular regulation. With a decade-long selection experiment on guppies *Poecilia reticulata*, we created populations with life histories adapted to positively size-dependent harvest, i.e., like that induced by fishing (fast life history). For comparison, we also created populations adapted to the opposite size-selection, and populations experiencing no size-selection. Fish exposed to positively size-dependent harvest were bolder, more likely to feed, were more social/aggressive, and expressed less brain *avt (*arginine vasotocin) relative to those exposed to negatively size-dependent harvest. In addition, higher expression of *th* and *th2* (tyrosine hydroxylase 1 and 2), and *neuroD2* (neuronal differentiation factor 2) were linked with bolder behaviour and higher feeding in normal (no-threat) conditions, while higher *avt*, *th*, and *neuroD2* were associated with higher sociability/aggression after a threat. Fish exposed to positively size-dependent harvest presented behaviours linked to faster life histories as theoretically expected. Therefore, harvest selection does not only affect fish size and life history, but indirectly leads to boldness and higher feeding rates, which potentially results in higher vulnerability to predators and higher pressure on prey abundance, respectively. Our results suggest that size-dependent mortality have further consequences to the ecosystem, beyond the target species.

## Introduction

Anthropogenic stressors are important drivers of ecological and evolutionary change. The impact of harvesting, i.e., humans acting as predators through fishing and hunting to exploit natural resources, is particularly large. This is due to the intense harvest rate imposed by humans compared to mortality imposed by natural predators and the breadth of the diversity affected by harvesting (Darimont et al. 2015, 2023; Jaureguiberry et al. 2022). As a result, harvesting is the most important driver of diversity loss in the marine environment, and the second most important on land (Di Minin et al. 2019). Harvesting does not only imply a large mortality rate relative to natural predators, but the mortality is disproportionately imposed on large individuals, due to preference, size-limit regulations, or economic value, resulting in selection opposite to that impose by natural selection (Diaz Pauli & Festa-Bianchet 2024; Heino et al. 2015).

Positively size-dependent mortality, such as that imposed by fishing, is expected to lead to evolutionary change towards smaller body size and faster life history (early maturation and increased fecundity), as has been observed in several marine species (Fugère & Hendry 2018; Heino et al. 2015). Moreover, small body size is likely correlated with behavioural traits that influence the energy flow in the ecosystem. These effects are felt both up and down the food webs. For example, increased consumption rate affects effectivity as a predator, and increased boldness leads to increased vulnerability to natural predators (Réale et al. 2010; Woodward et al. 2005). However, little is known on how size-dependent fishing can affect these behaviours and thus its potential impact on the ecosystems. Models assessing the effect of size-dependent fishing concluded that a bolder behaviour would be favoured in addition to fast life history (Andersen et al. 2018; Holt & Jørgensen 2015). These models also predicted an increase in boldness even when fishing was size-independent (Andersen et al. 2018; Holt & Jørgensen 2015). The few studies that have evaluated such indirect effects on behaviour concluded that fish exposed to positively size-dependent mortality became shyer and less likely to feed, which was opposite to the theoretical expectations (Diaz Pauli et al. 2020; Evangelista et al. 2021; Uusi-Heikkilä et al. 2015; Walsh et al. 2006). Another study showed that the behavioural response was sex-specific, with females becoming less likely to feed, but males becoming bolder (Diaz Pauli et al. 2019). These size-selection studies were performed under experimental conditions where harvesting occurred at a fixed age and density, and feeding was strictly controlled, avoiding the natural development of hierarchies within the populations. Therefore, earlier studies lacked density-dependent processes and thus ecologically relevant dynamics.

Here, we take advantage of a decade-long size-selection experiment on guppies, *Poecilia reticulata*, to evaluate the indirect effect of size-selection on ecologically relevant behaviours. In addition, to obtain an integrative understanding, we measured the brain expression of mRNA of candidate genes to study the neurobiological mechanisms driving the behaviours (Aubin-Horth 2016). Our laboratory selection experiment was designed to reach a higher degree of ecological realism than earlier studies, as populations were age-and size-structured, and density-dependent processes as well as “natural” and sexual selection were allowed (Bartusevičiūtė et al. 2022; Diaz Pauli 2012; Diaz Pauli et al. 2017). Our populations represent three different size-dependent harvest regimes: 1) Positively size-dependent harvest, where individuals above a minimum size-limit were removed, mimicking fishing-like selection, 2) Negatively size-dependent harvest, representing the opposite size-dependent selection, and 3) Size-independent harvest, where both large and small individuals were removed from the populations. Accordingly, the experimental populations represent three different life-history strategies. Populations exposed to positively size-dependent harvest were smaller in length, matured earlier, had increased fecundity and longer lifespan, relative to populations exposed to negatively size-dependent harvest. Populations exposed to size-independent harvest presented intermediate values between the two extremes. These changes reflect an evolutionary response to size-dependent harvest as they had a genetic basis (Diaz Pauli et al. unpublished).

We sampled individuals from our experimental guppy populations and evaluated feeding rate, boldness, and sociability, both under normal experimental conditions and after a threat. All three behaviours are ecologically relevant and linked to potential effects up and down the food web, as feeding rate can directly affect prey abundance, boldness determines encounter rate with predators, and sociability accounts for interaction with conspecifics and, in turn, both foraging and predator avoidance (Jolles et al. 2020; Moran et al. 2017; Réale et al. 2007). In addition, we quantified brain mRNA expression of several genes known to be linked to behavioural differences in fish. These included arginine vasotocin (*avt*; arginine vasopressin is the mammalian homolog), which is a neuropeptide that regulates a wide range of social behaviours, from vocal communication, sexual behaviour, aggression, parental behavioural to social recognition (Goodson & Bass 2001); tyrosine hydroxylase (*th*), and tyrosine hydroxylase 2 (*th2*) are involved in the synthesis of dopamine and other catecholamines and have been linked to anxiety-and stress-like behaviours, activity and other social behaviours (Kawabata et al. 2012); neuronal differentiation factors 1 and 2 (*neuroD1*, *neuroD2*) are involved in neural differentiation and hence neurogenesis and are highly expressed in shy and non-aggressive fish relative to bold fish (Johansen et al. 2012; Sørensen et al. 2013). We expected that fish exposed to positively size-dependent harvest would be bolder, less social, and more willing to feed, due to their faster life-history strategy (Réale et al. 2010; Woodward et al. 2005). Moreover, we expected fish exposed to positively size-dependent harvest to present lower expression of *neuroD* and *avt*, and higher levels of *th*, as such brain gene expression was observed in other fish species with similar behavioural patterns as described above (Castanheira et al. 2017; Reddon et al. 2022).

## Material and methods

### Experimental fish

Our experimental fish (N=108) originate from nine populations exposed to size-dependent harvest for over a decade. The three harvest regimes, each replicated thrice, were: 1) positively size-dependent harvest (hereafter referred as Positive harvest), which mimics selection on size as that imposed by common fishing; 2) negatively size-dependent harvest (referred as Negative harvest), representing the opposite size-selection than fishing and like the size-selection imposed by natural predators; and 3) size-independent harvest (referred as Random harvest). Details on the experimental design can be found in Bartusevičiūtė et al. (2022), Diaz Pauli (2012), and Diaz Pauli et al. (unpublished). The size threshold for harvesting was 16 mm (standard length, SL) since male guppies mature around this size (Reznick, Rodd, et al., 1996) and it is similar to minimum size limits typically imposed by fishing regulations. Harvesting started in 2010 and was performed every 6-12 weeks, with an average intensity of 40% (maximum 60% and minimum of 25%; Diaz Pauli et al. unpublished).

In June 2021, the first batch of 36 individuals were sampled from the populations (1:1 sex ratio, 12 fish/harvest regime) to assess their behavioural traits and the associated brain gene expression. The second batch of 72 fish (1:1 sex ratio, 24 fish/harvest regime) were sampled in September 2021 and were also assessed for behavioural traits and brain gene expression.

### Behavioural traits assessment

Experimental fish were placed in isolation in 3-litre tanks in a room that ensured water temperature at 26 ± 1°C and a photoperiod of 12:12 h (L:D). Fish were allowed to acclimatise to the new environment overnight and were fasted for 24 h before behavioural test. During the behavioural test, fish could not see each other to avoid learning from the experience of the neighbour fish. However, between the tests, visual interaction was allowed among the neighbours to ensure welfare.

Behavioural observations were performed on each fish under normal conditions and threatening conditions, immediately after a simulated predatory attack (being netted out of the tank for 2 seconds). Each of these two conditions was repeated twice. Therefore, each fish was exposed to 4 behavioural observations with a time lag of 2-3 days between each observation. The repetition of the same test occurred 5 days after the first observation, which allowed us to estimate short-term behavioural repeatability. We avoided a longer lag between repetitions as we wanted to assess the phenotype of the fish from the populations, rather than the fish acclimated to isolation for too long.

Behavioural observations started after providing the test fish with 3 drops of a solution of *Artemia salina* nauplii in water (x□ ± SD = 6.8 ± 3.5 *Artemia*/drop). The observation period lasted for 5 minutes. Three behavioural traits were measured: 1) Freezing time, as the time in seconds that the fish spent immobile in the tank, which is a proxy of shyness. 2) Feeding rate, as the number of bites the fish performed to ingest an *Artemia* nauplius, and 3) Time looking at their reflection on the tank wall, which is a proxy for social interactions with a conspecific. This measurement is similar to the mirror test, which measures social interactions in *Poecilia reticulata* and other fishes, as fish do not recognise their image and consider the reflection a conspecific (Castanheira et al. 2017; Savaşçı et al. 2021). However, the mirror test can measure social interactions from aggression to schooling, depending on experimental specifications and species (Savaşçı et al. 2021; Scherer et al. 2016). For female guppies, time looking to their own reflection is likely a prediction of sociability as using live conspecifics (Cattelan et al. 2017), while in males contact with the mirror might reflect aggressiveness (Budaev 1997).

### Gene expression analyses

One day after the last behavioural observation all fish were anaesthetised with buffered metacaine (0.3 g L^-1^) and then euthanised with an overdose (0.6 g L^-1^). The brain was dissected out and immediately placed in RNAlater, to preserve the integrity of the RNA (Ambion, USA) and stored for 24 h at 4°C and then at −80°C until subsequent mRNA expression analyses of *avt*, *neuroD1*, *neuroD2* and *neuroD2-like*, *th*, and *th2* using specific primers (Table 1). Total RNA was isolated from the guppy whole brain using TRI Reagent (Sigma-Aldrich, USA) following the manufacturer’s instructions. Quantity and integrity of total RNA was determined using a Qubit® 3.0 Fluorometer the Qubit® RNA BR (Broad Range) Assay Kit (ThermoFisher Scientific Inc., USA) and an Agilent 2100 Bioanalyzer (Agilent Technologies, USA), respectively. RNA integrity numbers (RIN) averaged 9.03 (0.55 SD). Afterwards, QuantiTect Reverse Transcription Kit (Qiagen, Germany) was used to remove possible genomic DNA contamination and synthesize cDNA from 250 ng of total RNA according to manufacturer’s protocol.

**Table 1.**
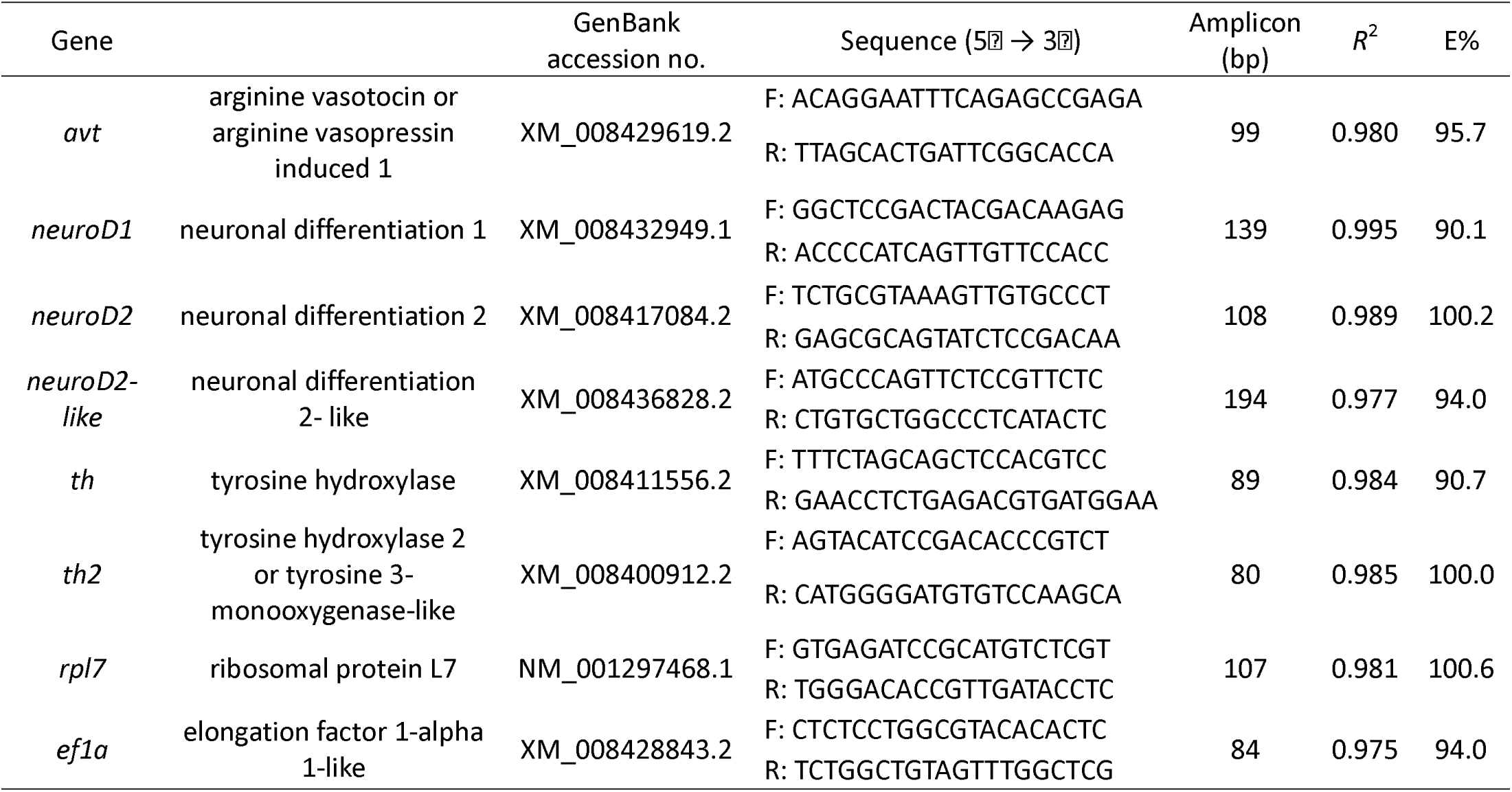
Sequences of the specific primers used for qPCR mRNA expression analysis. Primer sequences, amplicon sizes (bp), *R*^2^, and qPCR efficiency (E in %) are indicated for each primer pair.

The qPCR assay efficiency (*E*) was determined using a 5-point, 10-fold stepwise dilution series generated from a cDNA pool for each target gene and for the two reference genes, in triplicates. Melting curve analysis was performed over a range of 65–95°C (increment of 0.5°C for 0.05 s). *E* was calculated with the formula *E* = 10 ^(-1/slope)^ (Pfaffl 2001) and efficiencies ranged from 90 to 100.6% (Table 1). Gene transcription was analysed in a Bio-Rad CFX96 Touch Real-Time PCR system (Bio-Rad, USA) using iTaq™ Universal SYBR® Green Supermix (Bio-Rad, USA) in a total reaction volume of 12.5 μL per well, including 2.5 μL of 1:10 diluted cDNA and 0.25 µl of forward and reverse primers (10 μM). The cycling conditions were the following: 2 min at 95 °C followed by 37 cycles at 95 °C for 15 s and 60 °C for 25 s. Samples were run in duplicate. Relative transcription quantification was calculated using the mean normalized expression method and the geometric mean of the reference genes *rpl7* and ef1a (Vandesompele et al. 2002).

### Statistical analyses

All statistical analyses were performed with R (version 4.1.2; R Core Team 2021). Generalised linear mixed effect models were performed with glmmTMB package (version 1.1.5; Brooks et al. 2017) and diagnosed with DHARMa package (version 0.4.5; Hartig 2022). Behavioural repeatability were estimated with MCMCglmm package (version 2.34; Hadfield 2010). In addition, emmeans package (version 1.8.1-1; Lenth 2022) was used for pairwise comparisons.

All behavioural variables that were counts were zero-inflated (i.e., > 21% of data points were zero) and therefore analysed with statistical models that included zero-inflation. Furthermore, because of overdispersion, we assumed negative binomial distribution (rather than Poisson distribution). Freezing time is a real-valued variable, but it was converted to a binary response (0: zero seconds of freezing, 1: >zero seconds of freezing) and modelled with binomial distribution, because in the negative binomial zero-inflated model with real-valued response residual quantile deviations from uniformity were detected. All full models included fish ID nested within the two measurement repetitions as random effect, and fish length (standardised by subtracting 16 mm SL), threat treatment (No threat vs. After threat), harvest regime (Positive, Negative, Random), sex, and all the 2-way interactions as fixed effects. Models were simplified according to Akaike’s Information Criterion (AIC, Burnham et al. 2010). In addition, to assess the effect of gene expression on the different behaviours, similar models were performed for each behavioural variable where the full model included fish length, sex, and the expression of one gene at the time as fixed effects and population as random effect. This was done separately for two subsets of data, one including only the behaviours measure the first-time fish were tested under no threatening conditions, and the other including behaviours measured the first-time fish were tested after a threat.

Equivalent models were used to estimate behavioural repeatability, but using zero-inflated and Poisson distribution, as the MCMCglmm package does not include negative binomial distribution and already controls for overdispersion (Hadfield 2010). We estimated repeatability among measurements (n = 4) adjusted for experimental conditions (No-threat vs. After a threat) following Dingemanse & Dochtermann (2013). Repeatability and Bayesian credibility intervals (BCI) were estimated using a Bayesian approach with uninformative priors and the default settings of the library MCMCglmm. Statistical support for repeatability was determined by differences in Deviance Information Criteria (ΔDIC) between the models that included and excluded fish id as random effect (Dingemanse & Dochtermann 2013; Spiegelhalter et al. 2002).

Gene expression variables were transformed using natural logarithms and models with normal distribution were used. Full models included length standardised by harvest size threshold, sex, harvest regime and their 2-way interactions as fixed effects, while population was included as random effect. As above, models were simplified according to AIC.

## Results

### Behaviour

The three behavioural variables had high adjusted repeatability. Freezing time had the highest repeatability, R_a_ = 0.60 (95% CI=0.44–0.98; ΔDIC=-121), followed by Time looking reflection R_a_ = 0.55 (95% CI=0.42–0.72; ΔDIC=-46). Feeding rate had the lowest repeatability, R_a_ = 0.49 (95% CI=0.41–0.65; ΔDIC=-88). Table S1 shows between-and within-individual variances underlying these estimates.

Freezing time, analysed as a binary response (freezing or not), was affected by threat treatment (no-threat vs. after-threat), harvest regime, and their interactions (Table 2). The largest single effect was caused by the threat treatment (Figure 1; Table 2; Table S2): individuals under no threat were overall shyer, as they were highly likely to freeze, whereas exposure to threat greatly reduced the odds of freezing. Length and sex only affected freezing after the threat, where odds of freezing were reduced in smaller individuals and in males compared to females. Under no-threat conditions, individuals in random-harvest and positive-harvest treatments had reduced odds of freezing compared to the negative-harvest treatment fish. In the positive-harvest treatment, this difference was annulled after the threat.

**Figure 1.**
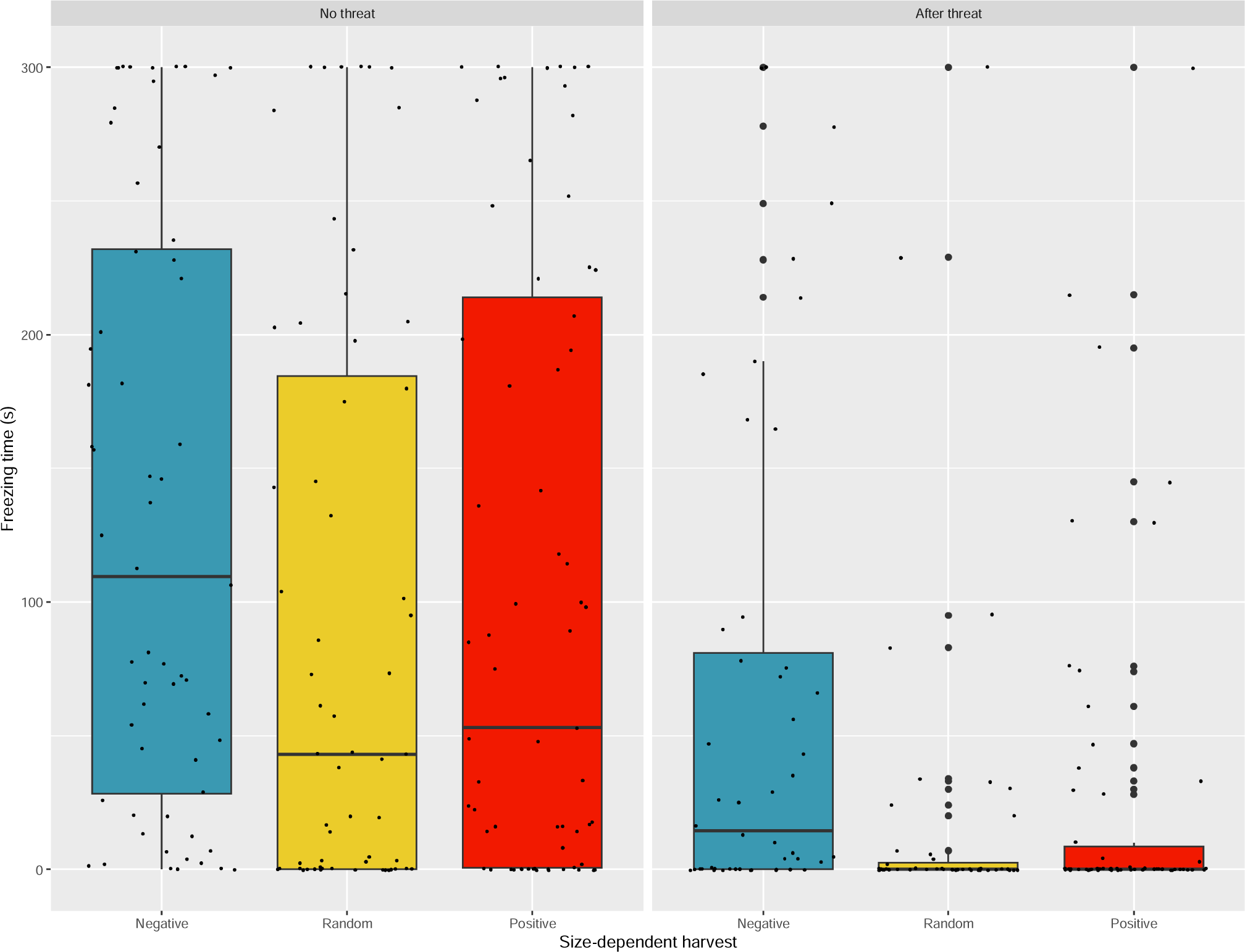
Time freezing for each harvest regime (blue: negative harvest, yellow: random harvest, red: positive harvest) in no-threat (left panel) and after threat (right panel) conditions. Small jittered black dots represent data points, while large dots represent outliers.

**Table 2.** Freezing time analysed as probability of freezing. Results from a generalized linear mixed effect model with binomial error distribution. The final model includes the categorical effects of threat condition (No-threat or After threat), size-dependent harvest (Negative, Random, or Positive), and sex (Female or Male), as well as standardized length as the regressor. Estimated coefficients are given as odds ratio and log-odds, with the associated confidence intervals (CI), *z* scores, and *p* values.

| Predictor | Odds Ratio | Log-Odds Estimate | Log-Odds CI | $z$ | $p$ |
| --- | --- | --- | --- | --- | --- |
| Intercept<br>(*Negative, No-threat, Female, 16 mm*) | $4.72 \times 10^{10}$ | 24.58 | 12.49 ... 36.67 | 3.98 | **0.0001** |
| Length | 0.48 | -0.74 | -1.93 ... 0.46 | -1.21 | 0.2269 |
| After threat | $2.7 \times 10^{-9}$ | -19.73 | -30.31 ... -9.16 | -3.66 | **0.0003** |
| Random | 0.00045 | -7.7 | -15.00 ... -0.40 | -2.07 | **0.0388** |
| Positive | 0.0001 | -9.24 | -16.72 ... -1.76 | -2.42 | **0.0154** |
| Male | 0.0016 | -6.45 | -14.20 ... 1.31 | -1.63 | 0.1033 |
| Length * After threat | 4.81 | 1.57 | 0.27 ... 2.87 | 2.36 | **0.0181** |
| After threat * Random | 167.2 | 5.12 | -1.17 ... 11.41 | 1.6 | 0.1105 |
| After threat* Positive | 21882 | 9.99 | 1.55 ... 18.44 | 2.32 | **0.0204** |
| After threat* Male | 0.00023 | -8.38 | -13.57 ... -3.20 | -3.17 | **0.0015** |

Feeding rate was affected both by threat treatment and by harvest regime. However, only the propensity to feed or not (zero-inflated model, Table 3) was affected, whereas the individuals that did feed had a similar number of bites across the treatments (count model, Table 3). Fish under no-threat conditions were less likely to feed on *Artemia* – i.e., they had 5 times higher odds of not feeding relative to fish after a threat (Table 3). In addition, harvest regime affected feeding rate in interaction with body length (Figure 2; Table 3; Table S2). At the standard-length threshold for harvesting (16 mm), fish exposed to negatively size-dependent harvest fed less – i.e., they had 3.9 times higher odds of not feeding – relative to positively size-dependent harvest (Figure 2b; odds ratio SE = 2.12 df = 326, *t*-ratio = 2.55, *p* = 0.0113). However, in fish exposed to positively size-dependent harvest, feeding rate decreased with increase in length (Increase in 1 unit of standard deviation, SD: odds ratio = 0.63, SE = 0.11, df = 326, *t*-ratio = -2.61, *p* = 0.0096). While in fish exposed to negative and random harvest feeding rate did not change with body length (Table 3). This led to smaller differences in the probability of feeding between negative and positive harvest at larger length (e.g., median length, 18 mm: odds ratio = 2.44, SE = 0.96, df = 326, *t*-ratio = 2.26, *p* = 0.0246), but larger differences between random and positive harvest (e.g., median length: odds ratio = 0.47, SE = 0.17, df = 326, *t*-ratio = -2.06, *p* = 0.04). And resulted in fish exposed to random harvest feeding more than the other harvest regimes at lengths above the average length (x□ ± SD = 19 ± 2.9 mm). Fish exposed to negatively size-dependent harvest consistently fed the least (Figure 2).

**Figure 2.**
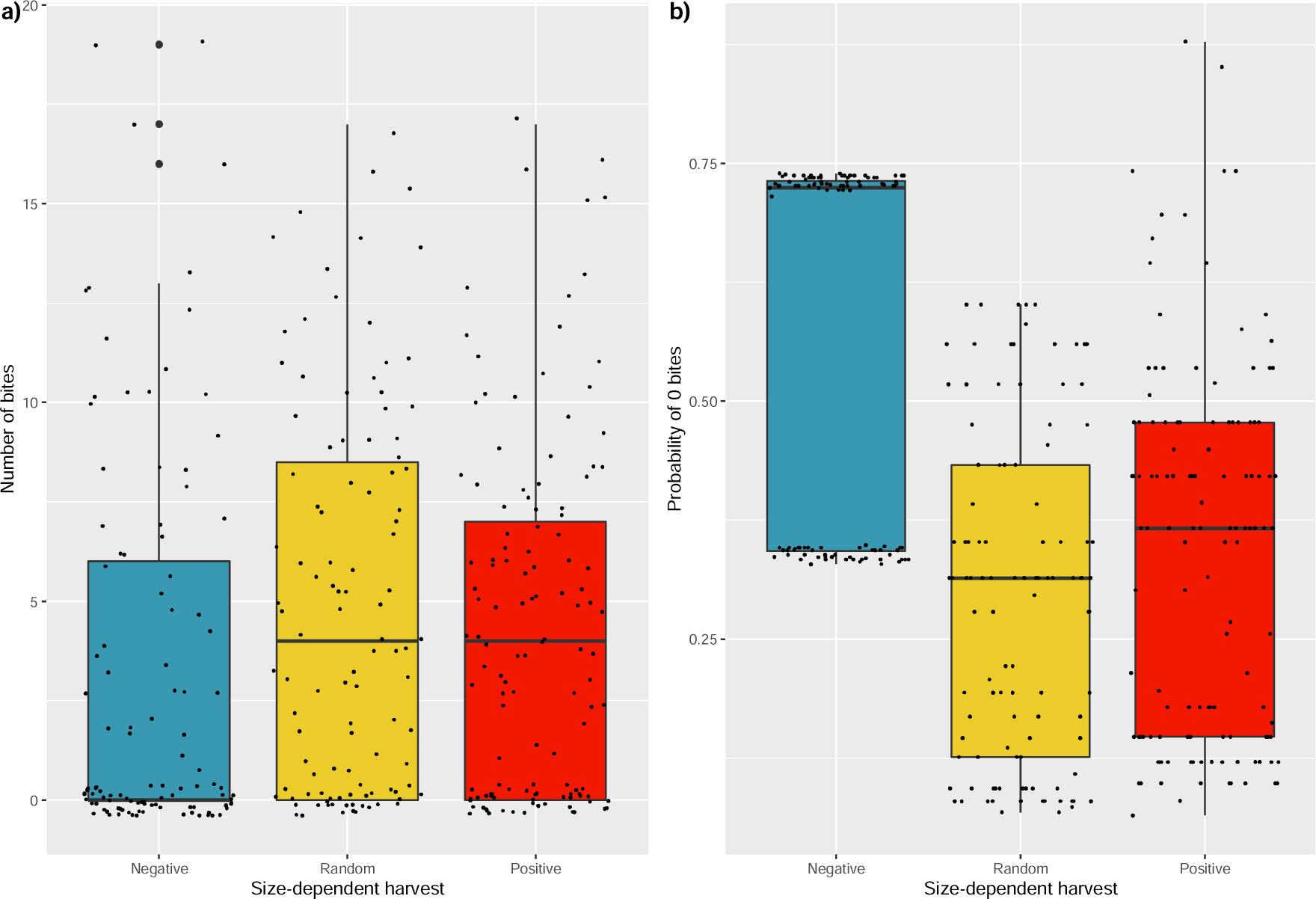
a) Feeding rate (number of bites) and b) predicted probability of taking zero bites for each harvest regime (blue: negative harvest, yellow: random harvest, red: positive harvest). Small jittered black dots represent data points in a) and model predicted fitted values in b).

**Table 3.** Effects of test conditions, size-dependent harvest, and sex on feeding rate. Output from generalized linear mixed effect model with zero-inflated negative binomial distribution. Estimated coefficients given in mean number of bites and logged-mean (count model), and in odds ratio and logged-odds (zero-inflated model), confidence intervals (CI), *z* and *p* values. Negative, Random and Positive refer to size-dependent harvest. Length refers to standardized length (see the methods). No-threat and After a threat refer to experimental test conditions, respectively.

| Count Model |  |  |  |  |  |  |
| --- | --- | --- | --- | --- | --- | --- |
| Predictors | Mean (bites) | Log-Mean | Log-CI | $z$ | $p$ | |
| Intercept |  |  |  |  |  |  |
| *(Negative, No-threat, Length=16 mm)* | 6.69 | 1.9 | 1.71 – 2.09 | 19.24 | **<0.001** |  |
| After threat | 0.97 | -0.03 | -0.27 – 0.20 | -0.27 | 0.787 |  |
| Length | 1.04 | 0.04 | -0.01 – 0.08 | 1.47 | 0.142 |  |

| Length * After threat | 0.95 | -0.06 | 0.12 – 0.005 | -1.8 | 0.072 |
| --- | --- | --- | --- | --- | --- |
| Zero-Inflated Model |  |  |  |  |  |
| Predictors | Odds ratio | Log-odds | Log-CI | z | p |
| Intercept |  |  |  |  |  |
| (Negative, No-threat, Length=16 mm) | 2.87 | 1.05 | 0.09 – 2.02 | 2.14 | **0.033** |
| After threat | 0.19 | -1.67 | -2.20 – -1.13 | -6.09 | **<0.001** |
| Length | 0.99 | -0.01 | -0.18 – 0.16 | -0.13 | 0.9 |
| Random | 0.13 | -2 | -3.25 – -0.75 | -3.14 | **0.002** |
| Positive | 0.25 | -1.37 | -2.43 – -0.32 | -2.55 | **0.011** |
| Length* Random | 1.2 | 0.18 | -0.06 – 0.43 | 1.45 | 0.148 |
| Length * Positive | 1.27 | 0.24 | -0.00 – 0.48 | 1.93 | 0.054 |

Time looking own reflections was affected by all experimental variables. The propensity of looking at own reflection or not was affected by length, sex, and harvest treatment (zero-inflated model, Table 4), whereas time looking among the individuals that did look (count model, Table 4) was affected by the said variables as well as by the threat treatment. Sex was the strongest effect in the propensity of looking at own reflection (zero-inflated model, Table 4; Figure S2a). Males were consistently more likely to look at their reflection relative to females (i.e, less probability of no looking; Female/Male: odds ratio = 3.05, SE = 1.62, df = 317, *t*-ratio = 2.10, *p* = 0.036). Harvest regime was the strongest single effect on the duration of time looking, but this effect depended on sex and length (count model, Table 4; Figure 3). At the size threshold, males exposed to positively size-dependent harvest looked the longest at their reflection (x□ ± SE 21 ± 6.3 s), while those exposed to negative harvest looked the shortest (x□ ± SE 0.54 ± 0.52 s) and those exposed to random harvest lay in between (Negative/Random: ratio = 0.13, SE = 0.12, df = 317, *t*-ratio = -2.17, *p* = 0.031; Negative/Positive: ratio = 0.025, SE = 0.03, df = 317, *t*-ratio = -3.61, *p* = 0.0004). For females, only those exposed to Positive harvest differed from females exposed to Negative harvest (Negative/Positive: ratio = 0.0005, SE = 0.001, df = 317, *t*-ratio = -3.49, *p* = 0.0005). At average population length, the differences among harvest regimes disappeared in males, while females exposed to Negative harvest spent less time looking than the other harvest regimes. For instance, averaged-sized females exposed to Positive harvest looked at their reflection 29 seconds more than averaged-sized females exposed to Negative harvest (Negative/Positive: odds ratio = 0.006, SE = 0.01, df = 317, *t*-ratio = -3.54, *p* = 0.0005). In addition, threat treatment also affected the duration of time looking at own reflection in interaction with sex and length (count model, Table 4; Figure S2b). Large individuals looked at their reflection more than short individuals (Table 4). However, this positive correlation between body length and time looking at reflection was steepest in males after the threat (100 s more/+1SD length increase; ratio = 0.02, SE = 0.017, df = 317, *t*-ratio = -4.86, *p* < 0.0001), and non-existent in females under no-threat conditions (1 s more/+1SD length increase; ratio = 0.85, SE = 0.39, df = 317, *t*-ratio = -0.36, *p* = 0.718). Therefore, at the size threshold, both males and females spent 4 seconds longer time looking at reflection in normal relative to threatening conditions (e.g., males: ratio = 2.79, SE = 0.91, df = 317, *t*-ratio = 3.14, *p* = 0.002). However, at mean population length (19 mm), there was no longer a difference between no-threat and threat conditions for males, while the 4 second difference remained for females (males: ratio = 0.57, SE = 0.23, df = 317, *t*-ratio = -1.41, *p* = 0.159; females: ratio = 3.62, SE = 1.54, df = 317, *t*-ratio = 3.03, *p* = 0.003).

**Figure 3.**
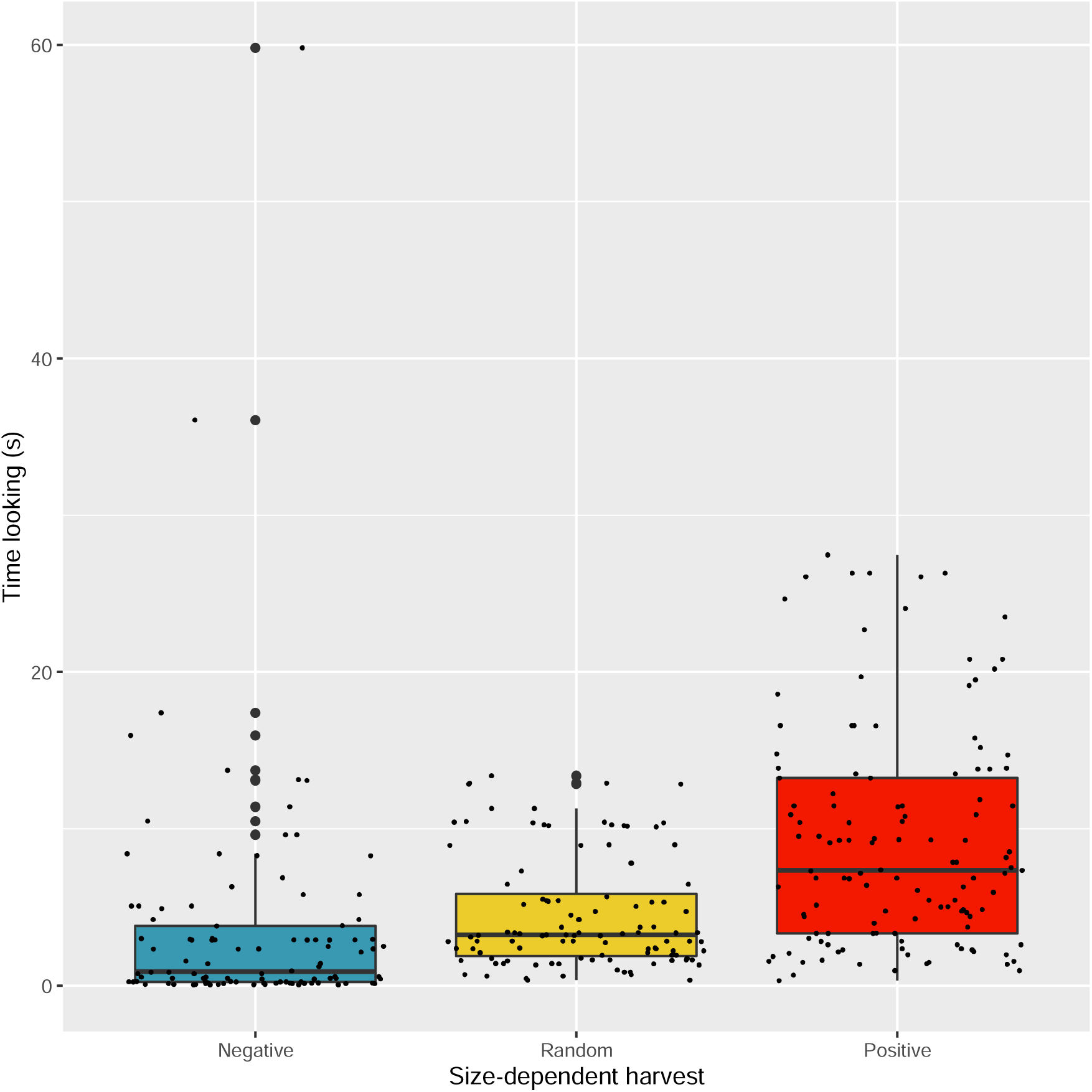
Predicted time looking at own reflection for each harvest regime (blue: negative harvest, yellow: random harvest, red: positive harvest). Small jittered black dots represent model predicted fitted values, while large dots represent outliers.

**Table 4.** Effects of test conditions, size-dependent harvest, and sex effects on time looking at reflection (in seconds). Output from generalized linear mixed effect model with zero-inflated negative binomial distribution. Estimated coefficients given in mean number of bites and logged-mean (count model), and in odds ratio and logged-odds (zero-inflated model), confidence intervals (CI), *z* and *p* values. Negative, Random and Positive refer to size-dependent harvest. Length refers to standardized length (see the methods). No-threat and After threat refer to experimental test conditions, respectively.

| Count Model |  |  |  |  |  |  |
| --- | --- | --- | --- | --- | --- | --- |
| Predictors |  | Mean<br>(seconds) | Log-Mean | Log-CI | *z* | *p* |
| Intercept<br>(*Negative, No-threat, Female, Length=16 mm*) |  | 0.08 | -2.58 | -6.68 – 1.52 | -1.23 | 0.218 |
| Length |  | 1.57 | 0.45 | 0.02 – 0.89 | 2.03 | **0.042** |
| Random |  | 109.96 | 4.7 | -1.01 – 10.41 | 1.61 | 0.107 |
| Positive |  | 2177.13 | 7.69 | 3.37 – 12.00 | 3.49 | **<0.001** |
| Male |  | 11.91 | 2.48 | -1.09 – 6.04 | 1.36 | 0.173 |
| After threat |  | 0.06 | -2.87 | -4.25 – -1.49 | -4.07 | **<0.001** |
| Length * Random |  | 0.68 | -0.39 | -1.14 – 0.37 | -1.01 | 0.314 |
| Length * Positive |  | 0.45 | -0.8 | -1.39 – -0.22 | -2.68 | **0.007** |
| Length * Male |  | 2.15 | 0.76 | 0.22 – 1.31 | 2.76 | **0.006** |
| Length * After threat |  | 1.67 | 0.51 | 0.25 – 0.77 | 3.85 | **<0.001** |
| Random * Male |  | 0.07 | -2.68 | -7.21 – 1.86 | -1.16 | 0.247 |
| Positive * Male |  | 0.02 | -4.02 | -7.76 – -0.27 | -2.1 | **0.035** |
| Male * After threat |  | 6.33 | 1.84 | 0.55 – 3.14 | 2.8 | **0.005** |
| Zero-Inflated Model |  |  |  |  |  |  |
| Predictors |  | Odds<br>ratio | Log-odds | Log-CI | *z* | *p* |
| Intercept<br>(*Negative, No-threat, Female, Length=16 mm*) |  | 18.01 | 2.89 | 1.04 – 4.75 | 3.05 | **0.002** |
| Length |  | 0.86 | -0.15 | -0.35 – 0.06 | -1.42 | 0.156 |
| Male |  | 0.08 | -2.58 | -4.72 – -0.43 | -2.35 | **0.019** |
| Random |  | 0.56 | -0.58 | -1.88 – 0.72 | -0.87 | 0.383 |
| Positive |  | 0.29 | -1.23 | -2.74 – 0.28 | -1.59 | 0.112 |
| Length * Male |  | 1.53 | 0.43 | 0.04 – 0.82 | 2.14 | **0.033** |
| Male * Random |  | 1.79 | 0.58 | -1.04 – 2.21 | 0.7 | 0.483 |
| Male* Positive |  | 6.92 | 1.93 | 0.00 – 3.86 | 1.96 | **0.05** |

### Behavioural and gene expression correlations

Behavioural differences are linked with dissimilar gene expression in the brain, but these depended on whether the behaviour was measured under no-threat or after threat conditions. Under no-threat, decreasing levels of *th2* mRNA expression were associated with 4.5 times higher odds of freezing (log-odds = -1.51, SE= 0.63, *N* = 98, *z* = -2.39, *p* = 0.017). Thus, individuals with high *th2* expression are bolder than individuals with low levels. Also under no-threat conditions, fish with lower mRNA expression of *th* and *th2* presented 4.9 and 3.7 times, respectively, the odds of not feeding (*th*: log-odds = -1.58, SE= 0.51, *N* = 98, *z* = -3.12, *p* = 0.0018; *th2:* log-odds = -1.31, SE= 0.53, *N* = 98, *z* = -2.46, *p* = 0.014). Thus, fish with higher *th* and *th2* expression were more likely to feed on *Artemia*. In addition, increasing mRNA expression of both *neuroD2* genes also resulted in higher probability of feeding on *Artemia* under no-threat conditions (*neuroD2*: log-odds = -1.09, SE= 0.51, *N* = 98, *z* = -2.16, *p* = 0.0307; *neuroD2-like:* log-odds = -1.16, SE= 0.59, *N* = 98, *z* = -1.98, *p* = 0.048). After a threat these relationships were no longer significant. However, after a threat, higher mRNA expression of both *avt* and *th* were linked to longer time looking at own reflection, being the effect of *avt* larger (*avt*: log-estimate = 9.13, SE= 2.25, *N* = 71, *z* = 4.06, *p* = 0.00005; *th*: log-estimate = 0.67, SE= 0.29, *N* = 70, *z* = 2.32, *p* = 0.020). In addition, increasing expression of *neuroD2* resulted in higher probability of looking at reflection after a threat (*neuroD2-like:* log-odds = -1.49, SE= 0.74, *N* = 70, *z* = -1.99, *p* = 0.046).

Despite the several links between gene expression and behaviour, a direct effect of harvest regimes on the gene expression was only found for *avt.* Females exposed to the different harvest regimes did not differ in *avt* gene expression (Figure 4; Table 5). However, males exposed to Positive harvest presented 1.7 and 1.9 times lower *avt* expression, relative to males exposed to Random and Negative harvest, respectively (Positive/Random: ratio = 0.55, SE = 0.11, df = 93, *t*-ratio = -3.01, *p* = 0.0034; Positive/Negative: ratio = 0.66, SE = 0.14, df = 93, *t*-ratio = -2.039, *p* = 0.044). These differences were not due to differences in body length as this was controlled for (Table 5).

**Figure 4.**
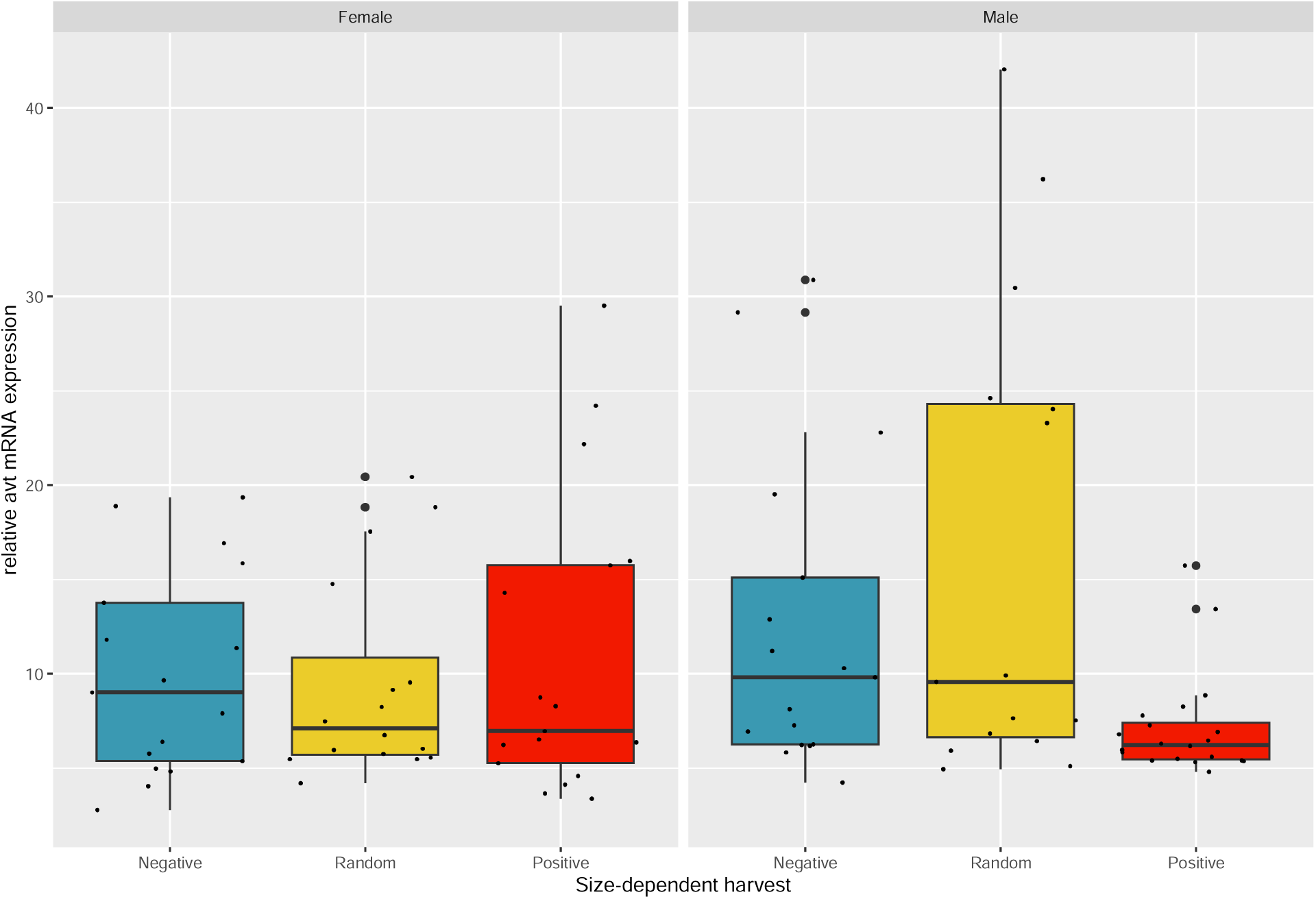
*avt* mRNA expression relative to reference genes (*rpl7* and *ef1a*) for each harvest regime (blue: negative harvest, yellow: random harvest, red: positive harvest) in females (left panel) and males (right panel). Small black dots represent data points, while large dots represent outliers.

**Table 5.** Effects of size-dependent harvest and sex effects on *avt* mRNA expression (log-transformed). Output from linear mixed effect model with gaussian distribution. Estimated coefficients given in relative expression, confidence intervals (CI), *z* and *p* values. Negative, Random and Positive refer to size-dependent harvest. Length refers to standardized length (see the methods).

| Predictors | Estimate | log-Estimate | CI | *z* | *p* |
| --- | --- | --- | --- | --- | --- |
| Intercept<br>(Positive, Female, Length=16 mm) | 8.58 | 2.15 | 1.85 – 2.46 | 13.89 | **<0.0001** |
| Length | 1.00 | 0.004 | -0.06 – 0.06 | 0.13 | 0.8973 |
| Random | 0.94 | -0.06 | -0.50 – 0.39 | -0.26 | 0.7972 |
| Negative | 0.96 | -0.04 | -0.51 – 0.43 | -0.15 | 0.878 |
| Male | 0.79 | -0.24 | -0.63 – 0.16 | -1.18 | 0.2374 |
| Random * Male | 1.92 | 0.65 | 0.09 – 1.21 | 2.26 | **0.0238** |
| Negative * Male | 1.58 | 0.46 | -0.08 – 1.00 | 1.66 | 0.0961 |

## Discussion

Fish exposed to positively size-dependent harvest for a decade, i.e., size-selection similar to that imposed by fishing, presented faster life histories as an evolutionary response (Diaz Pauli et al. unpublished) and were bolder, more likely to feed, looked more often at their reflection, and expressed less *avt* in the brain. In addition, we showed that higher mRNA expression of *th2* and *th* in the brain was linked to bolder behaviour and higher feeding rate under normal (i.e., no-threat) conditions, while higher brain expression of *avt* and *th* was linked to longer time looking to reflection after a threat. Here, we showed that size-dependent harvest indirectly leads to bolder and higher feeding rate. Therefore, size-dependent harvest does not only affect size and life-history traits but also ecologically relevant behaviours that can trickle up and down the food web, potentially affecting whole ecosystems.

Our harvest regimes were exclusively size-dependent, removing similar proportions of fish in all populations. The positively size-dependent harvest removed individuals larger than 16 mm, while the negatively size-dependent harvest removed individuals smaller than 16 mm. The former size-dependent harvest over a decade led to faster life history, while the latter resulted in slower life history (Bartusevičiūtė et al. 2022; Diaz Pauli et al. unpublished).

Here we showed that the populations further differed in behavioural traits. In general, fish exposed to positively size-dependent harvest became bolder, were more likely to feed, and spent longer time interacting with their own reflection, relative to fish exposed to negatively size-dependent harvest. Size-independent harvest also led to bolder, more frequent feeder and higher interaction with reflection, relative to negative harvest. It should be noted that behavioural differences were not due to size differences among populations (also seen in Harris et al. 2010), as these were controlled statistically. Therefore, our results agree with general evolutionary theory and specific size-dependent fishing models (Andersen et al. 2018; Holt & Jørgensen 2015; Réale et al. 2010; Woodward et al. 2005). However, earlier studies based on similar selection experiments concluded that fish exposed to fishing-like selection resulted in maladapted individuals, which were less willing to forage (Evangelista et al. 2021; Walsh et al. 2006) and shyer (Diaz Pauli et al. 2020; Uusi-Heikkilä et al. 2015; Walsh et al. 2006).

Differences among studies might be due to the different protocols used in assessing behaviours and to species-specific idiosyncrasies. However, difference in the design of the size-selection experiments themselves might be more important: earlier studies were performed using discrete-generations design where density was controlled and age diversity was minimal, thus limiting hierarchies and competition. Whether a behaviour is maladaptive obviously depends on the environment, as a shy behaviour and low willingness to forage might reduce per capita rate of energy gain and thus fitness in some contexts (Walsh et al. 2006), but not in some others: the opposite, bold behaviour and eager foraging, might increase encounters with predators (Holt & Jørgensen 2015; Nonacs & Blumstein 2010) and fishing gear (Arlinghaus et al. 2016; Diaz Pauli & Sih 2017). In general terms, shy and low-feeding fish might be better adapted to unstable and new environments, while bold and frequent feeders could be rewarded in stable environments (Castanheira et al. 2017). Therefore, for ecological resilience, maintenance of diversity might be the key, be it behavioural diversity or diversity in any other trait (Mimura et al. 2017; Smith & Blumstein 2013).

Repeatability estimates for the behaviours assessed here were above the average behavioural repeatability as estimated in meta analyses (Bell et al. 2009; Holtmann et al. 2017), indicating behavioural consistency in time and contexts (Réale et al. 2007). Moreover, high repeatability estimates indicate that the behaviours are stable and set an upper limit for heritability (Dochtermann et al. 2015; Killen et al. 2016). Therefore, the behaviours assessed here can experience evolutionary change. In addition, the behaviours are ecologically relevant, as they can directly influence energy flows up and down the food web (Jolles et al. 2020; Moran et al. 2017; Réale et al. 2007). For instance, lower feeding probability established in the lab in another small freshwater fish, *Oryzias latipes* (Diaz Pauli et al. 2019), was associated with lower benthic and planktonic prey abundance when the fish were raised in a mesocosm (Diaz Pauli et al. 2020; Evangelista et al. 2021). Therefore, behavioural differences found in the lab can be extrapolated to more realistic settings, and therefore, if present in the wild, can have ecosystem impacts. Moreover, the observed behavioural differences may also result in dissimilar vulnerabilities to fishing gear, thereby impacting the profitability of the fishery. For instance, fish more likely to feed might be more vulnerable to angling, while bold individuals are more likely to escape a trawl (Arlinghaus et al. 2016; Diaz Pauli & Sih 2017).

To better understand the neurobiological mechanism behind these behaviours we quantified the mRNA expression of several key genes involved in behavioural responses. In normal unthreatened conditions, higher mRNA expression of *th2* was linked to bolder behaviour, while higher *th, th2*, and *neuroD2* (both genes) mRNA expression was linked to higher feeding. Similar results were found for zebrafish, *Danio rerio*, and rainbow trout, *Oncorhynchus mykiss*, even though a correlation between activity and *th* mRNA expression was not directly assessed (Höglund et al. 2017; Omar et al. 2023). In zebrafish, individuals that moved shorter distance also presented lower *th* mRNA expression in the brain (Omar et al. 2023). Rainbow trout individuals with high activity and feeding motivation (referred as proactive) presented higher dopamine signalling (Höglund et al. 2017), which is related to high *th* expression (Robinson et al. 2016). However, our results linking higher expression of *neuroD2* with high feeding rate seems to oppose earlier studies in teleost fishes. In rainbow trout, it is reactive individuals (less feeding and active) who present higher *neuroD1* expression, linked with higher neural plasticity and flexible foraging (Øverli & Sørensen 2016).

After threatening conditions, higher *avt*, *th*, and *neuroD2* led to higher time looking at self-reflection of guppies in the present study. Higher *avt* mRNA expression is associated with higher aggression and reduced social interactions in several fishes (Desjardins & Fernald 2010; Oliveira et al. 2016; Reddon et al. 2022), including guppies (Cabrera-Álvarez 2018). Therefore, this reinforces the idea that time looking at self-reflection is a measure of aggression rather than sociability in the present study. Moreover, males spent more time looking at their reflection than females, which is consistent with the aggression interpretation: males are known to be more aggressive during foraging and mate competition than females (Magurran & Seghers 1991; Price & Helen Rodd 2006). Alternatively, increased *avt* mRNA expression could be linked to higher reactivity to stress and hyperactivity as an anxiety-linked behaviour (Mehr 2022; Reddon et al. 2022). Johansen et al. (2012) found that proactive rainbow trout presented lower *neuroD1* after long-term stress, but no difference under short-term stress (similar to after threat conditions in our study), while they did not assess *neuroD2*.

Neuroendocrine–behavioural associations have been intensively studied in rainbow trout (Castanheira et al. 2017). Using these results as the baseline, we could speculate about the meaning of our results in guppies. Here, guppies exposed to positively size-dependent harvest were bold, aggressive, and frequently-feeding, similar to the proactive rainbow trout. However, our aggressive and frequently-feeding guppies presented mRNA expression linked to higher neurological flexibility, rather than to lower flexibility observed in rainbow trout. Our results showed that differences in behaviour were linked to different mRNA expression and to direct effects of harvest regimes on the behaviours tested. However, we did not find direct effect of harvest regimes on mRNA expression, with *avt* mRNA expression being the exception. Males exposed to positively size-dependent harvest presented lower *avt* mRNA expression relative to the other harvest regimes. This contrasts with the result that positive harvest regime led to increased time looking at reflection, which is linked to high *avt* mRNA expression, as discussed above. Therefore, these opposing results require further study, preferably assessing differential expression in specific brain regions. Our results add to the growing body of studies on neurohormonal mechanisms regulating fish behaviour, highlighting a need for an integrative understanding of behaviour (Aubin-Horth 2016).

We found that populations exposed to positively size-dependent harvest (i.e., fishing-like size-selection) presented higher boldness, willingness to forage, and tendency toward social (probably antagonistic) interactions. Therefore, harvest based on size alone can indirectly affect a suit of behavioural traits that may impact food webs. Our experimental size-selection, despite being based on a laboratory set-up, allowed higher ecological realism than earlier selection experiments (Diaz Pauli et al. unpublished) and led to a behavioural response according to expectations (Andersen et al. 2018; Holt & Jørgensen 2015; Réale et al. 2010; Woodward et al. 2005). Moreover, we present insights into the neurobiological mechanisms behind the behavioural response. Increased boldness might be linked to higher vulnerability to predators and hence higher natural mortality, while increased willingness to forage may lead to higher impact on prey abundance (Holt & Jørgensen 2015; Moran et al. 2017). Therefore, our results contribute to the growing evidence that harvesting impacts populations and ecosystems beyond the targeted traits and single-population effects (e.g., Diaz Pauli et al. 2020; Evangelista et al. 2021; Uusi-Heikkilä et al. 2015; Walsh et al. 2006; Wood et al. 2018). This implies that fisheries management should consider an ecosystem approach that incorporates the wider impacts of fishing, for the benefit of sustainability and for the fishery itself.

## Acknowledgements

We are grateful to Diep Mach Ellertsen and Vitalija Bartusevičiūtė for support in the lab, Lars Ebbeson and Naouel Gharbi for helpful discussion when planning the experiments, and the Research Council of Norway for funding (project number 275125). The experiments described followed the Norwegian and University of Bergen animal welfare regulations and were approved by Norwegian Food Safety Authority (FOTS id 21812, accepted 12.02.2020).

---

**Table S1.** Estimated between-and within-individual variance of the behavioural traits from generalised mixed effect models, with the 95% confidence intervals in parentheses.

|  | Between-individual variance | Within-individual variance |
| --- | --- | --- |
| Freezing time (seconds) | 6.12 (2.86, 48.12) | 0.17 (0.08, 37.43) |
| Bites (count) | 0.55 (0.44, 1.01) | 0.52 (0.40, 0.77) |
| Bites (probability) | 15.40 (4.27, 56.42) | 18.79 (18.80, 18.80) |
| Look refection (count) | 19.90 (12.12, 39.20) | 16.30 (10.23, 25.20) |
| Look refection (probability) | 49.58 (18.27, 846.91) | 213.124 (213.12 213.15) |

**Table S2.**
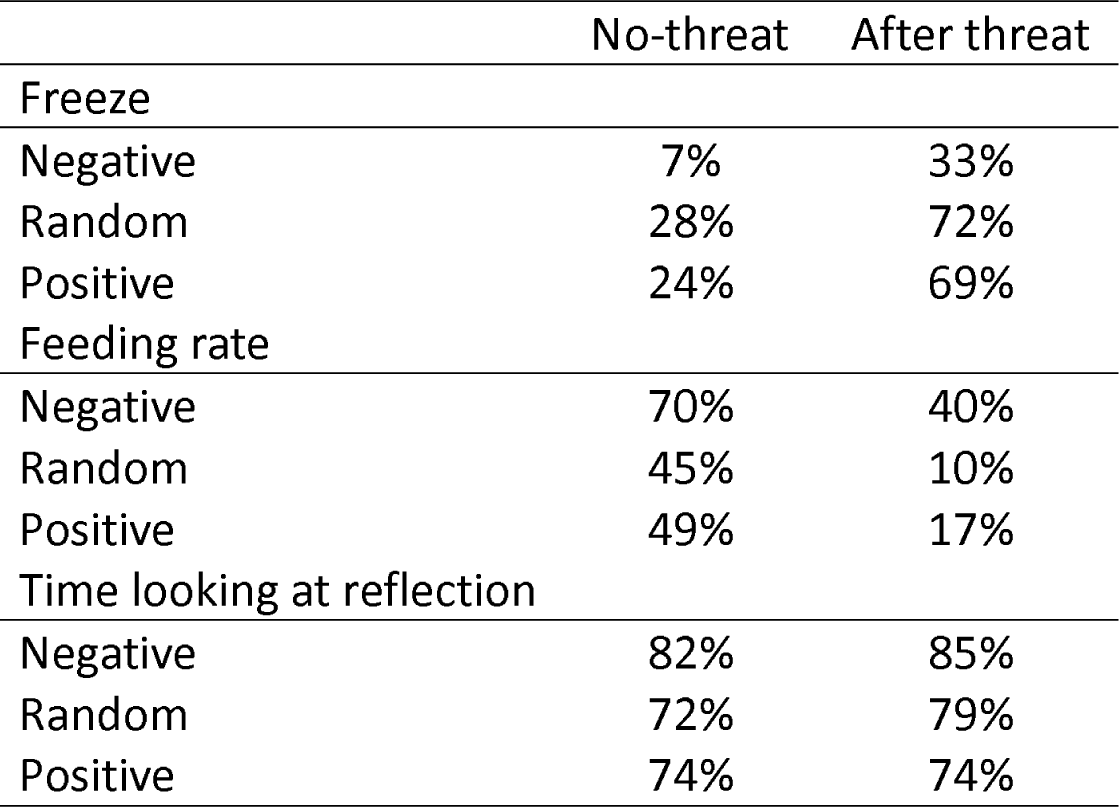
Proportion of zeros of Time freezing, Feeding rate, and Time looking at own reflection per harvest regime and threat treatment.

|  | No-threat | After threat |
| --- | --- | --- |
| Freeze |  |  |
| Negative | 7% | 33% |
| Random | 28% | 72% |
| Positive | 24% | 69% |
| Feeding rate |  |  |
| Negative | 70% | 40% |
| Random | 45% | 10% |
| Positive | 49% | 17% |
| Time looking at reflection |  |  |
| Negative | 82% | 85% |
| Random | 72% | 79% |
| Positive | 74% | 74% |

**Figure S1.**
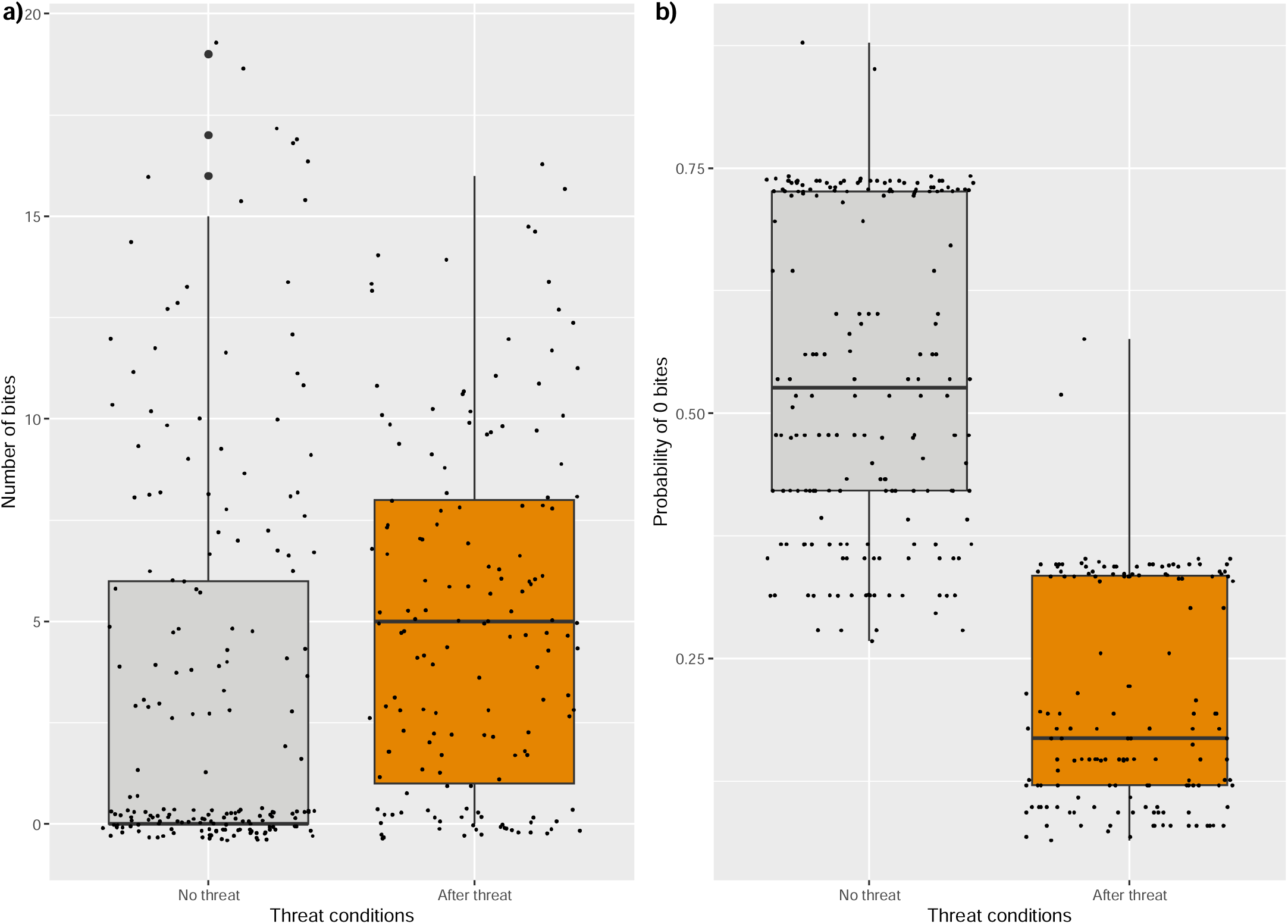
a) Feeding rate (number of bites, raw data) and b) predicted individual probability of taking zero bites for each threat level (grey: No threat, orange: after a threat). Small jittered black dots represent data points in a) and predicted fitted values in b).

**Figure S2.**
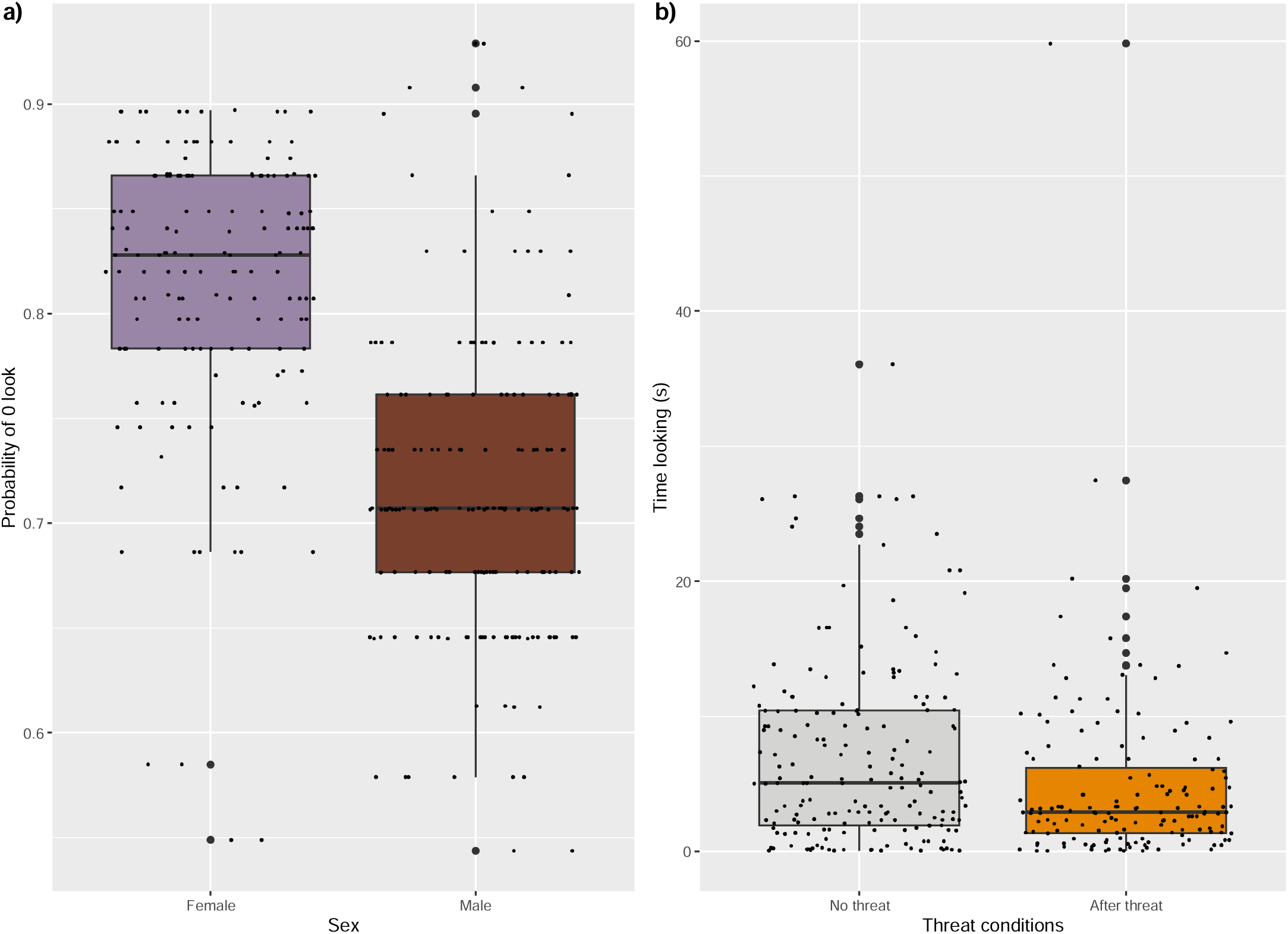
a) Predicted probability of looking zero seconds for females (purple) and males (brown), and b) predicted time looking (duration in seconds) for each threat level (grey: No threat, orange: after a threat). Small jittered black dots represent model predicted fitted values.

